# Takens’ theorem to assess EEG traces: regional variations in brain dynamics

**DOI:** 10.1101/2025.01.19.633767

**Authors:** Arturo Tozzi, Ksenija Jaušovec

**Author notes:** (corresponding author) 1155 Union Circle, #311427 Denton, TX 76203-5017 USA.

## Abstract

Takens’ theorem (TT) proves that the behaviour of a dynamical system can be effectively reconstructed within a multidimensional phase space. This offers a comprehensive framework for examining temporal dependencies, dimensional complexity and predictability of time series data. We applied TT to investigate the physiological regional differences in EEG brain dynamics of healthy subjects, focusing on three key channels: FP1 (frontal region), C3 (sensorimotor region), and O1 (occipital region). We provided a detailed reconstruction of phase spaces for each EEG channel using time-delay embedding. The reconstructed trajectories were quantified through measures of trajectory spread and average distance, offering insights into the temporal structure of brain activity that traditional linear methods struggle to capture. Variability and complexity were found to differ across the three regions, revealing notable regional variations. FP1 trajectories exhibited broader spreads, reflecting the dynamic complexity of frontal brain activity associated with higher cognitive functions. C3, involved in sensorimotor integration, displayed moderate variability, reflecting its functional role in coordinating sensory inputs and motor outputs. O1, responsible for visual processing, showed constrained and stable trajectories, consistent with repetitive and structured visual dynamics. These findings align with the functional specialization of different cortical areas, suggesting that the frontal, sensorimotor and occipital regions operate with autonomous temporal structures and nonlinear properties. This distinction may have significant implications for advancing our understanding of normal brain function and enhancing the development of brain-computer interfaces. In sum, we demonstrated the utility of TT in revealing regional variations in EEG traces, underscoring the value of nonlinear dynamics.

## INTRODUCTION

The human brain operates as a sophisticated nonlinear system, adept at handling extensive information via dynamic interactions (Khoshnoud et al., 2018; Zhao et al., 2020; Dai et al., 2022; Biloborodova et al., 2024). Electroencephalography (EEG) serves as a non-invasive, high-resolution method for investigating brain activity. Nonetheless, conventional linear analysis techniques frequently fall short in representing the intricate nonlinear features of EEG signals (Alturki et al., 2020). To address this limitation, nonlinear dynamics and chaos theory have emerged as powerful frameworks for understanding brain activity, with Takens’ theorem (henceforward TT) providing a cornerstone. TT establishes that the behavior of a dynamical system can be reconstructed in a multidimensional phase space using time-delayed versions of a single time series from the observed data (Takens 1981). In the context of EEG analysis, TT provides a robust mathematical tool to study temporal evolution, revealing properties that linear methods cannot uncover (Rohrbacker 2009). By reconstructing the phase space, researchers can analyze key EEG dynamical properties such as temporal dependencies, dimensional complexity and predictability (Kwessi and Edwards, 2021). This approach has proven valuable for identifying changes in neural dynamics associated with various cognitive and pathological conditions (Fell et al., 2000).

Previous research has highlighted the effectiveness of TT in analyzing EEG signals, especially for identifying pathological conditions like epilepsy, Alzheimer’s disease and schizophrenia (Kannathal et al., 2005; Altindiş et al., 2021; Cai et al., 2024; Al Fahoum and Zyout, 2024). However, less attention has been given to the application of this approach for assessing regional variations in brain dynamics under normal conditions. Different brain regions exhibit distinct patterns of electrical activity, reflecting their specialized roles in cognition, sensation and motor function. For instance, the frontal region (FP1) is associated with higher cognitive processes such as decision-making and working memory. The sensorimotor cortex (C3) governs movement and integrates sensory inputs, while the occipital region (O1) processes visual information. Despite their unique roles, interactions among these regions contribute to the brain’s global dynamics.

Comparing the spread and trajectory of phase space trajectories across different brain regions can shed light on how functional specialization translates into distinct dynamical features. In this study, we apply TT to assess EEG traces from three key brain regions: FP1, C3 and O1. We reconstruct phase spaces and analyze the temporal structure and complexity of the trajectories. Key metrics such as trajectory spread were computed to quantify regional differences in brain dynamics.

The rationale for using TT lies in the fact that overlapping trajectories reveal shared dynamical patterns across the EEG traces, suggesting common temporal structures in brain activity. These patterns likely arise from similar neural processes or consistent experimental conditions and reflect shared features of brain dynamics that remain consistent across individuals or trials. Conversely, differences in the spread or divergence of trajectories highlight variability in the dynamics of EEG signals, capturing subject-specific neural activity. This variability can serve as a marker for distinguishing between populations, such as healthy versus pathological groups, and is particularly relevant for identifying neural behaviors associated with specific cognitive states, tasks or conditions. Trajectories that densely fill the phase space indicate higher complexity, which is often linked to intricate neural processes and sophisticated information management. In contrast, smoother and more confined trajectories suggest simpler, more periodic or more deterministic behavior, highlighting a system with lower complexity but higher predictability. Together, these observations shed light on the dynamic interplay between stability, variability and complexity in brain activity, offering a deeper understanding of its temporal and spatial organization.

In the following sections, we detail the methods used for EEG data collection, phase space reconstruction and analysis, followed by the presentation and discussion of our results.

## SUBJECTS AND METHODS

This study investigates the regional variations in brain dynamics by applying Takens’ theorem to EEG data, focusing on three key channels: FP1 (frontal), C3 (sensorimotor), and O1 (occipital). The data consisted in the retrospective evaluation of ten EEG traces. EEG recordings were obtained from ten healthy, right-handed volunteers (mean age: 20.1 years; SD = 1.1; age range: 18–22 years; 5 males). For additional information on the participants and EEG methodologies, see Jaušovec and Jaušovec (2005) and Tozzi et al. (2021). The signals were captured using a 64-channel EEG system, adhering to the standard 10-20 electrode placement system. Preprocessing included artifact removal, such as eye blinks and muscle movements through independent component analysis and band-pass filtering (0.5–50 Hz).

After preprocessing, the analysis focused on reconstructing the phase space of each EEG trace. TT was employed to create a multidimensional representation of the system’s dynamics using delay embedding. The first step involved determining for each signal the optimal time delay (τ), a critical parameter in phase space reconstruction, as it defines the separation between consecutive points in the reconstructed dimensions (Matilla-García et al., 2021). The mutual information method was used to calculate τ, as it effectively identifies the point at which the time series exhibits the least redundancy while retaining information about the system’s dynamics. For all channels and traces, τ was consistently found to be 25, suggesting a shared temporal dependency structure across the EEG signals.

The next step was to determine the embedding dimension (d), which represents the number of dimensions required to unfold the systems dynamics without overlap in the reconstructed phase space (Xu 2018). The False Nearest Neighbors (FNN) method was used for this purpose (Rohrbacker 2009). By analyzing the proportion of neighbors that remain close when the dimensionality is increased, FNN identifies the point at which the embedding dimension captures the true dynamics of the system. The results consistently indicated d=6 for all traces, suggesting a moderate level of complexity in the underlying neural dynamics.

Starting from the parameters τ=25 and d=6, the phase space for each EEG trace was reconstructed and visualized in three dimensions by selecting the first three delay coordinates. The trajectories were analyzed to extract quantitative metrics characterizing the temporal structure of the phase spaces. Specifically, the spread of the trajectories in each dimension and the average distance between consecutive points were computed. The spread reflects the range of dynamical variability, while the average distance provides insight into the smoothness and temporal evolution of the signals.

### Tools and statistical analysis

Computational analysis for precise quantification of regional variations in brain dynamics was implemented using Python, leveraging libraries for signal processing, nonlinear analysis and statistical evaluation. Statistical analysis was performed to compare the phase space metrics among the FP1, C3 and O1 channels. Welch’s t-tests were used to evaluate differences in trajectory spread across dimensions (x, y, z) among the channels. Welch’s t-test was chosen because it does not assume equal variances between groups, thus ensuring robustness in the presence of heterogeneity in EEG signals.

## RESULTS

Phase space trajectories were reconstructed for all ten EEG traces of healthy individuals across the FP1, C3 and O1 channels using time delay τ=25 and embedding dimension d=6. These reconstructions highlighted distinct patterns of activity in the three regions (**Figure A**). To quantify these differences, key metrics such as trajectory spreads across dimensions (x, y, z) and average distances between consecutive points were computed for each channel (**Figure B**).

The FP1 trajectories displayed the broadest and most dispersed paths as well as the highest average distances, indicative of the greatest variability and complexity. This pattern aligns with the frontal region’s role in higher-order cognitive functions, such as decision-making and problem-solving, which demand flexible and dynamic neural processes. In contrast, the C3 and O1 trajectories were more compact. Among them, O1 exhibited the most constrained dynamics with the smallest spreads and average distances, reflecting the structured and repetitive nature of visual processing. This stability is characteristic of the occipital region’s specialization in handling consistent sensory inputs. Meanwhile, C3 showed a moderate spread, with intermediate spreads and distances that balanced variability and stability, consistent with its role in integrating sensory inputs and motor outputs, a process requiring balance between stability and flexibility. The degree of similarity and overlap in C3 trajectories also suggests shared neural patterns within the sensorimotor region, while the divergence in trajectory spreads provides insights into individual differences in neural activity among participants.

Statistical comparison of trajectory spreads across the three channels further validated these findings. Significant differences were observed in all pairwise comparisons, confirming substantial dynamical differences among FP1, C3 and O1. **Figure D** provides a direct comparison of trajectory spreads across dimensions (x, y, z) for the three channels. FP1 consistently showed the largest spreads across all dimensions, a finding that reflects the region’s high dynamical variability and complexity. This statistical validation reinforces the notion that EEG dynamics are regionally specific and closely linked to the functional roles of these cortical areas.

In sum, the combination of trajectory visualizations, quantitative metrics and statistical analysis reveal clear regional variations in trajectory spreads. These findings align with the functional specialization of the brain regions and underscore the utility of TT in capturing regional variations in EEG dynamics.

**Figure A.**
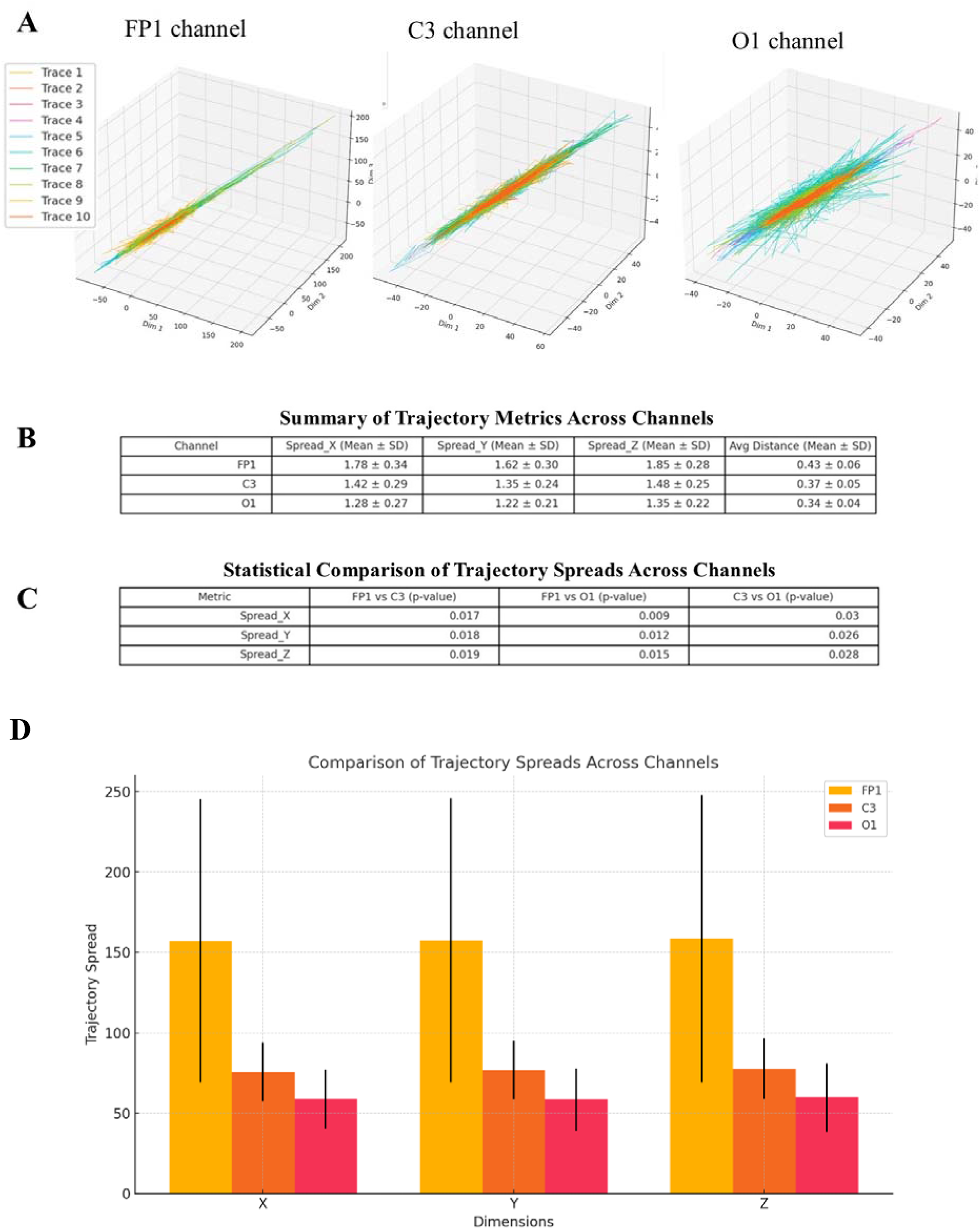
Comparison of trajectory spreads across the reconstructed phase spaces for the three EEG channels FP1, C3 and O1. Each phase space illustrates the combined trajectories of all ten EEG traces from the corresponding channel, visualized together in a single 3D plot. **Figure B**. Trajectory metrics computed for all three channels, including trajectory spreads across dimensions (x, y, z) and average distances between consecutive points. **Figure C**. Statistical comparison of trajectory spreads across the three channels. Welch’s t-tests reveals significant distinctions (p<0.05) across all pairwise comparisons. **Figure D**. Detailed visualization of the statistical comparison of trajectory spreads. The bar chart illustrates the average spreads for each dimension (x, y, z) for the FP1, C3 and O1 channels, accompanied by error bars representing the standard deviations.

## CONCLUSIONS

We focused on applying Takens’ theorem to EEG signals to examine regional nonlinear variations in healthy subjects’ brain dynamics, reconstructing the phase spaces and comparing their temporal features. Three key cortical areas were analyzed, namely, the frontal region (FP1), the sensorimotor region (C3) and the occipital region (O1). Using nonlinear dynamical analysis focused on trajectory spreads and average distances, the study reconstructed the phase space of EEG signals to reveal distinct patterns of activity corresponding to the functional specialization of these regions.

Takens’ theorem has emerged as a powerful tool for exploring EEG traces. By transforming abstract EEG data into comprehensible geometrical forms, TT allows for the reconstruction of the phase space. Rohrbacker (2009) illustrated the impact of TT on neuroscience by calculating the necessary embedding dimensions for EEG data using false nearest neighbor analysis. His seminal findings emphasized the importance of adequate embedding for accurately reconstructing phase spaces. TT’s applications span epilepsy diagnosis, causal inference, BCI development and real-time analysis. For instance, phase space analysis has been used to identify pre-seizure states by detecting chaotic patterns in EEG signals (Kwessi & Edwards, 2021). Kwessi and Edwards (2021) introduced the concept of complex geometric structurization to analyze epilepsy-related EEG data, revealing intricate geometric structures as biomarkers for seizure activity. Sleep studies have used phase space trajectories to differentiate between sleep stages and detect sleep disorders (Fell et al., 2000). Changes in neural complexity and stability underlying neurodevelopmental disorders such as ADHD can also be examined through phase space reconstruction (Kaur et al., 2020).

Topological data analysis (TDA) represents another area where TT has influenced EEG research (Xu et al., 2021). Altindiş et al. (2021) explored the use of TDA to extract persistent homologies from EEG-derived state spaces, demonstrating that TDA could robustly capture topological features of neural data even in the presence of artifacts. Moreover, TT has been instrumental in the development of methodologies for analyzing the nonlinear components of EEG signals. For instance, Mekler (2008) introduced an innovative method for calculating the correlation dimension of EEG attractors, overcoming the limitations of traditional approaches. This advancement allows for more efficient processing of large datasets while minimizing subjectivity. Similarly, Kannathal et al. (2005) applied entropy measures derived from EEG embeddings to detect epilepsy, achieving high classification accuracy. Also, TT has proven valuable for studying causality in neural systems. Traditional causal models often fall short when applied to nonlinear and cyclic systems like the brain. Harnack et al. (2017) addressed this limitation by leveraging time-delay state space reconstructions to measure directed causal influences. The applicability of TT extends to brain-computer interface (BCI) systems, which depend on decoding EEG signals to enhance signal classification and facilitate interaction with external devices, providing critical support for individuals with disabilities. Carrara and Papadopoulo (2024) incorporated embedding techniques into geometric neural networks, enhancing the interpretability and efficiency of BCI decoding.

The novelty of this study is that we applied TT to investigate the differences in phase space features among different EEG channels of healthy subjects, reflecting the distinct physiological dynamics of the underlying brain regions. One of the primary contributions of this work was the detailed reconstruction of phase spaces for each EEG channel using time-delay embedding. The reconstructed trajectories were further quantified through metrics such as trajectory spread and average distance that provided insights into the temporal structure and complexity of brain activity in different regions that are not easily captured by traditional linear methods. The FP1 region, associated with higher-order cognitive functions such as decision-making and information processing, exhibited the largest trajectory spreads and higher average distances. This variability aligns with the complexity and flexibility required for cognitive tasks. In contrast, the C3 region, which integrates sensory inputs and motor outputs, exhibited moderate variability, while the O1 region, responsible for visual processing, displayed the smallest spreads and distances, consistent with its structured and stable neural dynamics.

These findings have significant implications for understanding the role of regional dynamics in neural processing. The distinct temporal structures observed across the three channels provide a quantitative framework for examining how different brain regions contribute to overall function. By quantifying and visualizing the underlying dynamics, TT opens new opportunities for studying neural systems. The moderate embedding dimensions identified in this study suggest that the brain operates within a finite-dimensional dynamic space, efficiently processing complex information. The possible applications of these findings are vast. Dynamical systems can be compared by analyzing the variability, complexity and chaotic behavior of specific EEG traces. Identifying outliers, such as traces that deviate significantly from the shared structure, may help detect noise or reveal unique dynamics.

Despite its strengths, the study has limitations. The sample size was relatively small, consisting of only ten EEG traces, which may limit the generalizability of the findings. Additionally, the uniform use of τ=25 and =6 across all traces, while robust, may not capture individual variability in optimal parameters. Furthermore, the analysis was limited to three channels (FP1, C3, O1) representing specific cortical regions. Expanding the study to include more channels and larger datasets could provide a more comprehensive understanding of the global and regional dynamics of the brain. Future research could address these limitations by applying the approach to more diverse datasets, exploring alternative parameter optimization methods, and incorporating task-based EEG data.

In conclusion, this study demonstrates the utility of Takens’ theorem in reconstructing the phase space of EEG signals in healthy individuals and in revealing regional variations in brain dynamics. The findings highlight distinct patterns of activity in the frontal, sensorimotor and occipital regions. By bridging the gap between theoretical nonlinear dynamics and practical applications, this approach offers significant potential for advancing both basic neuroscience and clinical practice.

## DECLARATIONS

## Ethics approval and consent to participate

This research does not contain any studies with human participants or animals performed by the Authors.

## Consent for publication

The Authors transfer all copyright ownership, in the event the work is published. The undersigned authors warrant that the article is original, does not infringe on any copyright or other proprietary right of any third part, is not under consideration by another journal, and has not been previously published.

## Availability of data and materials

all data and materials generated or analyzed during this study are included in the manuscript. The Authors had full access to all the data in the study and take responsibility for the integrity of the data and the accuracy of the data analysis.

## Competing interests

The Authors do not have any known or potential conflict of interest including any financial, personal or other relationships with other people or organizations within three years of beginning the submitted work that could inappropriately influence, or be perceived to influence, their work.

## Funding

This research did not receive any specific grant from funding agencies in the public, commercial, or not-for-profit sectors.

## Authors’ contributions

The Authors equally contributed to: study concept and design, acquisition of data, analysis and interpretation of data, drafting of the manuscript, critical revision of the manuscript for important intellectual content, statistical analysis, obtained funding, administrative, technical, and material support, study supervision.

## Declaration of generative AI and AI-assisted technologies in the writing process

During the preparation of this work, the authors used ChatGPT to assist with data analysis and manuscript drafting. After using this tool, the authors reviewed and edited the content as needed and takes full responsibility for the content of the publication.

## Acknowledgements

none.

## Notes

### Competing Interest Statement

The authors have declared no competing interest.

